# Cell-to-cell and type-to-type heterogeneity of signaling networks: Insights from the crowd

**DOI:** 10.1101/2021.03.23.436603

**Authors:** Attila Gabor, Marco Tognetti, Alice Driessen, Jovan Tanevski, Baosen Guo, Wencai Cao, He Shen, Thomas Yu, Verena Chung, Single Cell Signaling in Breast Cancer DREAM Consortium members, Bernd Bodenmiller, Julio Saez-Rodriguez

**Author notes:** These authors contributed equally to the work (Attila Gabor, Marco Tognetti).

## Abstract

Recent technological developments allow us to measure the status of dozens of proteins in individual cells. This opens the way to understand the heterogeneity of complex multi-signaling networks across cells and cell-types, with important implications to understand and treat diseases such as cancer. These technologies are however limited to proteins for which antibodies are available and are fairly costly, making predictions of new markers and of existing markers under new conditions a valuable alternative. To assess our capacity to make such predictions and boost further methodological development, we organised the Single Cell Signaling in Breast Cancer DREAM challenge. We used a mass cytometry data set, covering 36 markers in over 4,000 conditions totalling 80 million single cells across 67 breast cancer cell lines. Through four increasingly difficult subchallenges, the participants predicted missing markers, new conditions, and the time course response of single cells to stimuli in the presence and absence of kinase inhibitors. The challenge results show that despite the stochastic nature of signal transduction in single cells, the signaling events are tightly controlled and machine learning methods can accurately predict new experimental data.

**Graphical Abstract:** 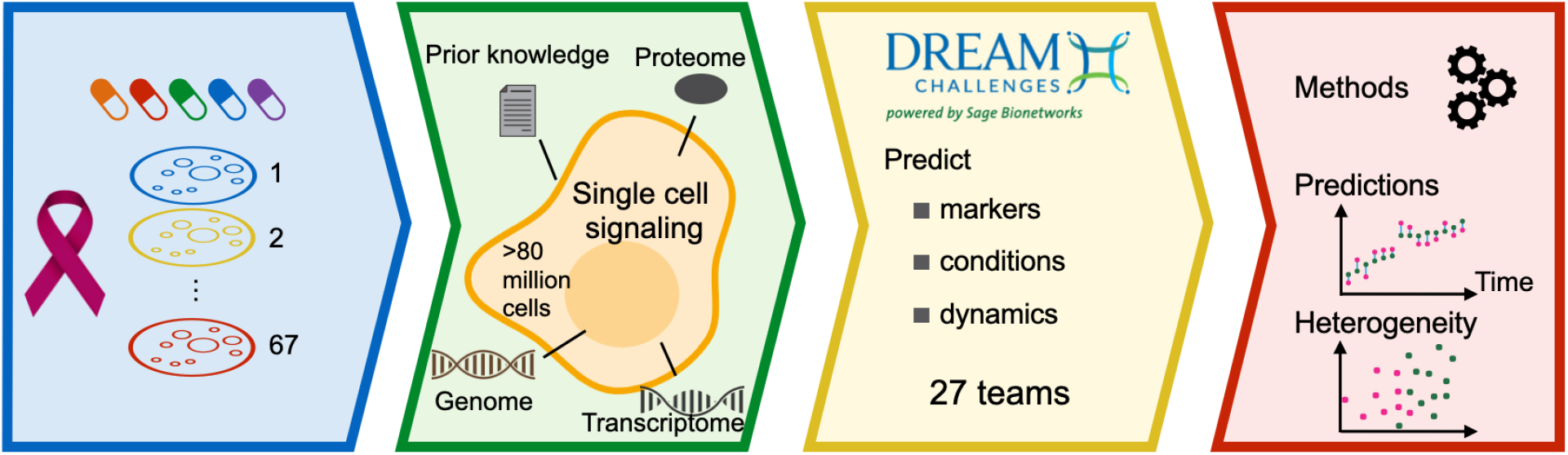

**Key points:**

- Over 80 million single-cell multiplexed measurements across 67 cell lines, 54 conditions and 10 time points to benchmark predictive models of single cell signaling
- 73 approaches from 27 teams for predicting response to kinase inhibitors on single cell level, and dynamic response from unperturbed basal omics data
- Predictions of single marker models correlate with measurements with a correlation coefficient of 0.76
- Top models of whole signaling response models perform almost as well as a biological replicate
- Cell-line specific variation in dynamics can be predicted from basal omics

## 1 Introduction

Cell signaling governs most activities of cells as it underlies the ability to correctly respond to stimuli from the cellular microenvironment. This dynamic process of signaling is tightly regulated by the interactions of proteins. Deregulation and major disturbances in this finely tuned machinery can lead to diseases such as cancer (Hynes and MacDonald 2009), autoimmunity (Benveniste et al. 2014), and diabetes (Boucher, Kleinridders, and Kahn 2014).

The abundance of signaling proteins and the mutation status of underlying genes vary across cell lines, therefore different cell lines have distinct signaling networks. But not only different cell lines have different signaling networks: cells of the same type display dissimilar signaling patterns in response to stimuli depending on history, cell state, and microenvironment. This heterogeneity has important implications for the treatment of diseases. For example, intratumor heterogeneity in cancer is a key contributor to therapeutic failure and drug resistance: often subpopulations do not respond to therapies, resulting in relapses. Thus a better understanding of signaling and its heterogeneity might unlock better treatments (Yaffe, 2019).

Single cell measurements, particularly mass cytometry (Bodenmiller et al. 2012), (Spitzer and Nolan 2016), have opened a new way to monitor the signaling in individual cells. With this technique we can measure the cellular response to multiple perturbations, paving the way for understanding the heterogeneity of the underlying cellular mechanisms.

However, the measured nodes are limited to a handful of proteins for fluorescence based live cell imaging and several dozens for highly multiplexed mass cytometry. If we could predict nodes that are not measured via computational approaches, we could overcome this technological limitation. In addition, the number of combinatorial treatments in studies is often limited by the budget or available material, since these experiments are complicated and expensive. Therefore, it would also be desirable if models could predict the single cell response of a cell line to new treatments after training on response data of other cell lines. One could even conceive models that predict the response to perturbations from only basal, unperturbed data.

Answering these questions requires advanced computational methods. Predictive mathematical models are often used for bulk data analysis (Rukhlenko et al. 2018), (Byrne et al. 2016), but they have not yet been applied extensively to single-cell data (Loos et al. 2018). Therefore, as a first step, we need to assess the available modeling frameworks for single cell predictions on a fair platform and explore their use, capabilities and also their limits.

To accelerate the development of methods, we organised the Single Cell Signaling in Breast Cancer (SCSBrC) DREAM challenge. The Dialogue for Reverse Engineering Assessments and Methods (DREAM) Challenges provides a framework for participants across the globe to compare their methods for solving biomedical problems (Saez-Rodriguez et al. 2016). The participants are ranked according to predefined metrics and the solutions are analysed to find the best ways to approach the particular problems. Further, the predictions, the methods and the code that reproduce the results are made publicly available after the challenge as a stepping stone for further improvements.

We based the SCSBrC challenge on a single-cell signaling mass cytometry dataset (Tognetti et al. 2020). In this dataset, 67 cell lines were stimulated with EGF in combination with one of five kinase inhibitors (PKC, PI3K, mTOR, MEK, EGFR), and in each condition 31 phosphoproteins and 5 cellular markers were measured at 10 different time points over the course of one hour (Figure 1, Supp. Figure 1 and Supp. Figure 2A). In total, this dataset includes more than 80 millions single cells and more than 4,000 experimental conditions. The data is complemented with population-level protein abundance experiments, transcriptomics data (RNA-seq) and genomics data (Single Nucleotide Polymorphism and Copy Number Aberration)(Marcotte et al. 2016). A portion of this data was kept for scoring (test data), and the rest was provided as training data to the participants to develop their methods. Further, we encouraged participants to incorporate external data and resources of prior knowledge, such as pathways and protein-protein interaction databases, in their models.

**Figure 1.**
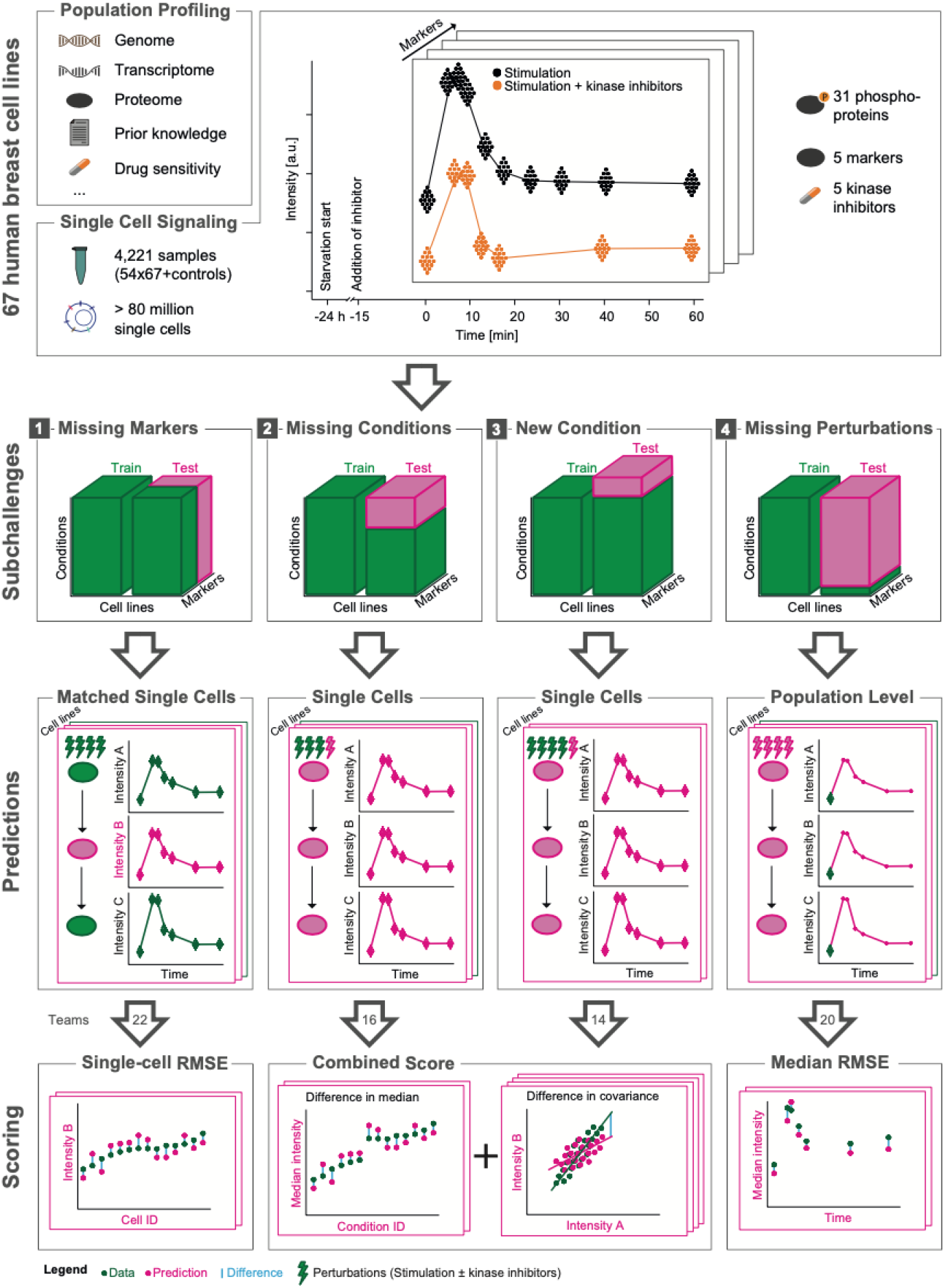
Overview of the Single Cell Signaling in Breast Cancer challenge. The challenge data consist of genomics, basal transcriptomics and proteomics data, and perturbation single cell phosphorylation datasets from 67 breast cancer cell lines. The datasets were divided into training (green; given to participants) and test (pink; withhold for scoring). The training data was common but test data different for each subchallenge. In each subchallenge a suitable metric was chosen to score the predictions.

Specifically, in our challenge we aimed to answer the following questions of increasing complexity: Can we predict the signaling response (1) of nodes that are not directly measured from other measurements on the same cells?, (2) to new combinatorial treatments based on how other cell lines respond to that treatment?, (3) to perturbations for which we have no data but know the target? and (4) purely from basal omics data? We defined four corresponding subchallenges (Figure 1).

We evaluated 73 models from 27 teams across the four subchallenges and found that machine learning based methods worked well for these prediction tasks. Subchallenge 1 showed that the signaling dynamics in nodes that are measured in some cell lines can be accurately predicted in other cell lines. In some cell lines the basal phosphorylation of the nodes is significantly different from others, which may lead to a shift between measurements and predictions, therefore we recommend to include the basal phosphorylation in the predictive models. Subchallenge 2 showed that the effect of kinase inhibitors can be learnt from other cell lines and predicted with high accuracy. Predictions performed better than random even when only the target of the kinase inhibitor is given (Subchallenge 3). Further, we found major differences between model types in terms of performance at capturing intra-cell line heterogeneity. Finally, we found in Subchallenge 4 that models can predict the dynamics of a cell line from the basal omics if one uses the general signaling response of other cell lines.

## 2 Results

The SCSBrC DREAM challenge was organised in three rounds between September and December 2019. A total of 261 individuals registered, and 22, 16, 14 and 20 teams around the world submitted predictions in the final round of each subchallenge, respectively. The challenges were followed by a post-challenge period where participants submitted write-ups and code, and participated in a survey about their methods. The experimental data, models and results are freely available for further use (https://www.synapse.org/singlecellproteomics).

### 2.1 The missing marker prediction subchallenge

The goal of subchallenge 1 (SC1) was to build a model that can predict the phosphorylation of a node that one may not be able to measure experimentally. To obtain such models, we asked participants to predict the phosphorylation of selected proteins in single cells from specific cell-lines and conditions (pink box, Figure 1). To do this, the participants could use the other nodes of the signaling network for the same condition and cell-lines as well as the same nodes from different cell-lines (green box; Figure 1; Supp. Figure 1). Specifically, we asked the participants to predict five markers (p-ERK^Thr202/Tyr-204^, p-AKT^Ser473^, p-PLCg2^Tyr759^, p-HER2^Tyr1196^, p-S6^Ser235/Ser236^) in six cell lines, measured in all the 49 different conditions, totalling 11.9 millions cells. The participants submitted their predictions for each measured single cell, and we compared them to the actual measurements of the markers using the root-mean-square error (RMSE) (see Methods and Figure 1). This performance was also compared against a reference model we built using a random forest predictor (see Methods).

#### 2.1.1 Quality of predictions

First, we ranked the teams and compared their performance with random predictions and with a random forest based reference (see Methods). For robust ranking, we used bootstrap sampling to generate a distribution of scores for each team (see Methods; Figure 2A). This identified the winner (*icx_bxai*, RMSE_icx_bxai_ = 0.849), which strongly outperformed other participants (Bayes factor > 499). For the random prediction, we took the test data and shuffled the unique cell-ids in each cell line and experimental condition. This way, when we scored this randomised dataset, each cell was predicted by a randomly chosen other cell in the same cell line and experimental condition. Almost all teams performed better than this random prediction (Figure 2A), with 32% average improvement (RMSE_teams average_ = 0.979 +-0.238, RMSE_random_= 1.44 +-2.6e-4). Six teams achieved better predictions than a reference model that we built before the challenge started (RMSE_reference_=0.903, see Methods).

**Figure 2:**
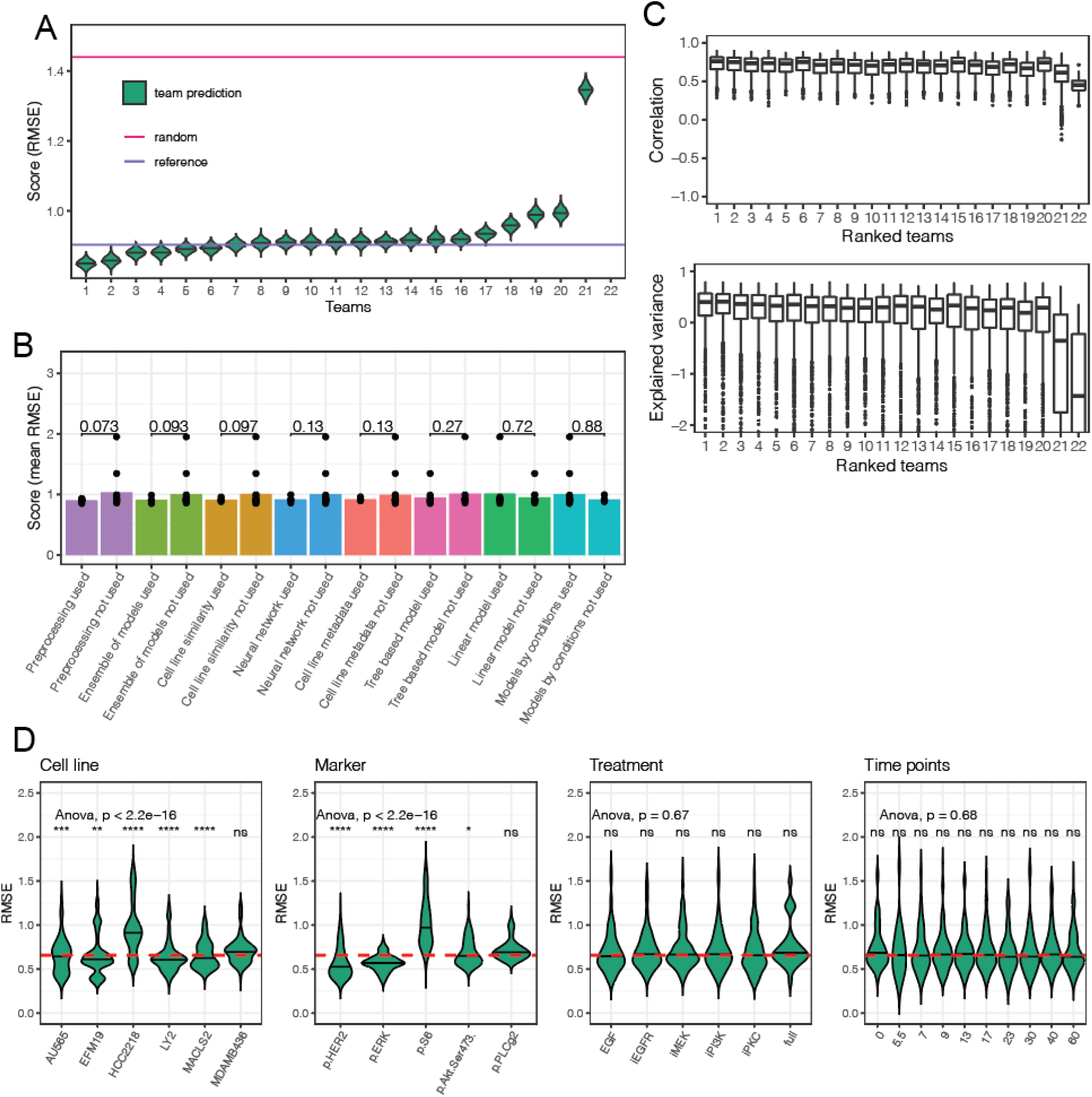
summary of subchallenge 1. A) rank of the teams in SC1 based on the RMSE. A random and the baseline prediction from our model that we built prior to the challenge (see Methods) are also shown for reference. B) the barplot compares the mean performance of the teams using a specific method or data set against the other teams; individual performances are marked with dots. Numbers above the bars represent the p-value from a t-test. C) Distribution of the correlation coefficients and explained variances across conditions, cell lines and markers, for each team. D) distribution of the prediction errors for cell lines, markers, treatments and time points. Each violin is built from the prediction errors of all teams. The dashed red horizontal line shows the global mean RMSE. The significance of a t-test between the groups and the global RMSE is shown above each violin plot (*: p<0.05, **: p<0.01, ***: p<0.001, ****: p<0.0001, ns: not significant)

Models predicted the trend of the data well, but for a few cell lines the predictions under- or overestimated the test data. Model predictions correlated strongly with the test data (median correlation coefficient: 0.761) in each condition (Figure 2C top) and explained large amounts of variance (R^2^_median_ = 0.40, R^2^_max_ = 0.803) across all teams (Figure 2C bottom). The explained variance was negative in some conditions, which means that the predictions of node phosphorylation were worse than the sample mean in those conditions. These conditions were found to belong to specific cell lines that had strongly different initial activation (at time 0, before stimulating) of the predicted nodes than the majority of cell lines (see section 2.1.3), leading to a shift between predictions and measurements.

The training data did not contain measurements of the initial activation of the test cell lines. To quantify how much the knowledge of this initial phosphorylation influences the accuracy of the predictions, we corrected the predictions by the difference between the predicted and measured mean basal activities of the cell lines. All the predictions improved significantly (Supp. Figure 3D): in average teams’ error decreased by 10.4 %, and even for the top team the error decreased by 7% (from RMSE_icx_bxai_=0.849 to RMSE_icx_bxai, corrected_=0.789). This suggests that it is very hard to predict phosphorylation without any data for the phospho-protein of interest in the cell-line where it has to be predicted.

#### 2.1.2 What makes a good model for node prediction?

To understand what makes a model better than others, we analyzed the participants’ write-up of methods and their answers to a questionnaire (Supp. File 1). We compared the building blocks of the methods to see which elements contributed to better predictions (Figure 2B). Although we did not find a single model type or modelling procedure strikingly beneficial, several features improved the accuracy of predictions. In particular, pre-processing of the data (mean RMSE improvement: 0.13, p.value = 0.0731 from a t-test), using an ensemble of models (estimated RMSE improvement: 0.094, p.value = 0.0935) and training models on similar cell lines (improvement: 0.0949, p.value = 0.0975) had the largest individual impact on the score.

While some teams relied on a single method to build their models, the top ranked team (*icx_bxai*) built a model for each marker independently and included the 32 provided markers, treatment and time, and the top principal components of the proteomics data as features. Further, when estimating the marker values at time t_i_, they also considered the median signaling levels of the 32 measured markers at time t_i_, t_i-1_ and t_i+1_. They used subsets of these features to train an ensemble of models: ElasticNet, ExtraTrees, RandomForest, Light Gradient Boosting and linear model. To make a prediction, they averaged the predictions of all models.

The successful combination of methods by the best performers suggest that, as experienced in other DREAM challenges (Saez-Rodriguez et al. 2016; Marbach et al. 2012), the combinations of methods provide robust predictors. We performed multiple aggregation of the predictions (see Methods and Supp. Figure 3C), but could not achieve significant improvement: the combination of the top two teams led to 0.5% improvement over the best team. This is probably due to the large correlation between the residuals of the teams predictions, which means that when a team overshot with their predictions, other teams did similarly.

#### 2.1.3 Time course analysis of predictions

We investigated if there were specific markers, conditions or cell lines that are harder to predict than others (Figure 2D). Prediction errors distributed heterogeneously across the cell-lines and markers (ANOVA test, p.value < 2.2e-16, Supp. Table 1), but homogeneously across treatments (p.value = 0.67) and time after the perturbation (p.value = 0.68) (Figure 2D). In particular, *p-Her2* was the most accurately predicted marker (improvement of 0.142 of the global mean RMSE, Supp. Table 2) and the largest prediction error was found for *p-S6* (deterioration of 0.305). Cell line HCC2218 has the largest prediction errors (0.246 above global mean RMSE), while other cell lines had similarly small prediction error values. The prediction error differences between cell lines and markers suggest that the cell lines show diverse signaling patterns that influence the models’ accuracy (Supp. Figure 4).

As signaling is a dynamic process, we aggregated the predictions of each team and the test data at each time point and compared the time-courses. The dynamic response of phosphosites changed strongly across the different perturbations and also between cell lines (Supp. Figure 4), which was captured well by the predictions. For example, MDAMB436 cells showed strong p-AKT activation upon all perturbations with EGF, except when a PI3K inhibitor (iPI3K) was applied. The majority of models captured both the initial dynamics and the level at which the phosphorylation saturated and predicted the low response of p-AKT when iPI3K was added. The measured members of the Mitogen-activated Protein Kinase (MAPK) pathway (e.g. p-MEK, p-ERK, and downstream p-p90RSK) show similar activation dynamics in response to EGF stimulations, which was accurately predicted by most participants (Figure 3B). For example, there is a strong decrease in p-ERK when we compare its response between EGF and iMEK conditions (mean phosphorylation decreases: log_2_ fold change -7.66, adj. p-value = 4.14e-23 (Figure 3A)), which was also predicted well by most models (Figure 3B). However, in five cell lines (four training cell lines: HCC2185, HCC1599, DU4475, CAL148 and one test cell line HCC2218) the p-ERK phosphorylation did not decrease after inhibiting MEK (Figure 3B), indicating a rare (observed for 5 out of 67 cell lines) MEK independent ERK signaling.

**Figure 3:**
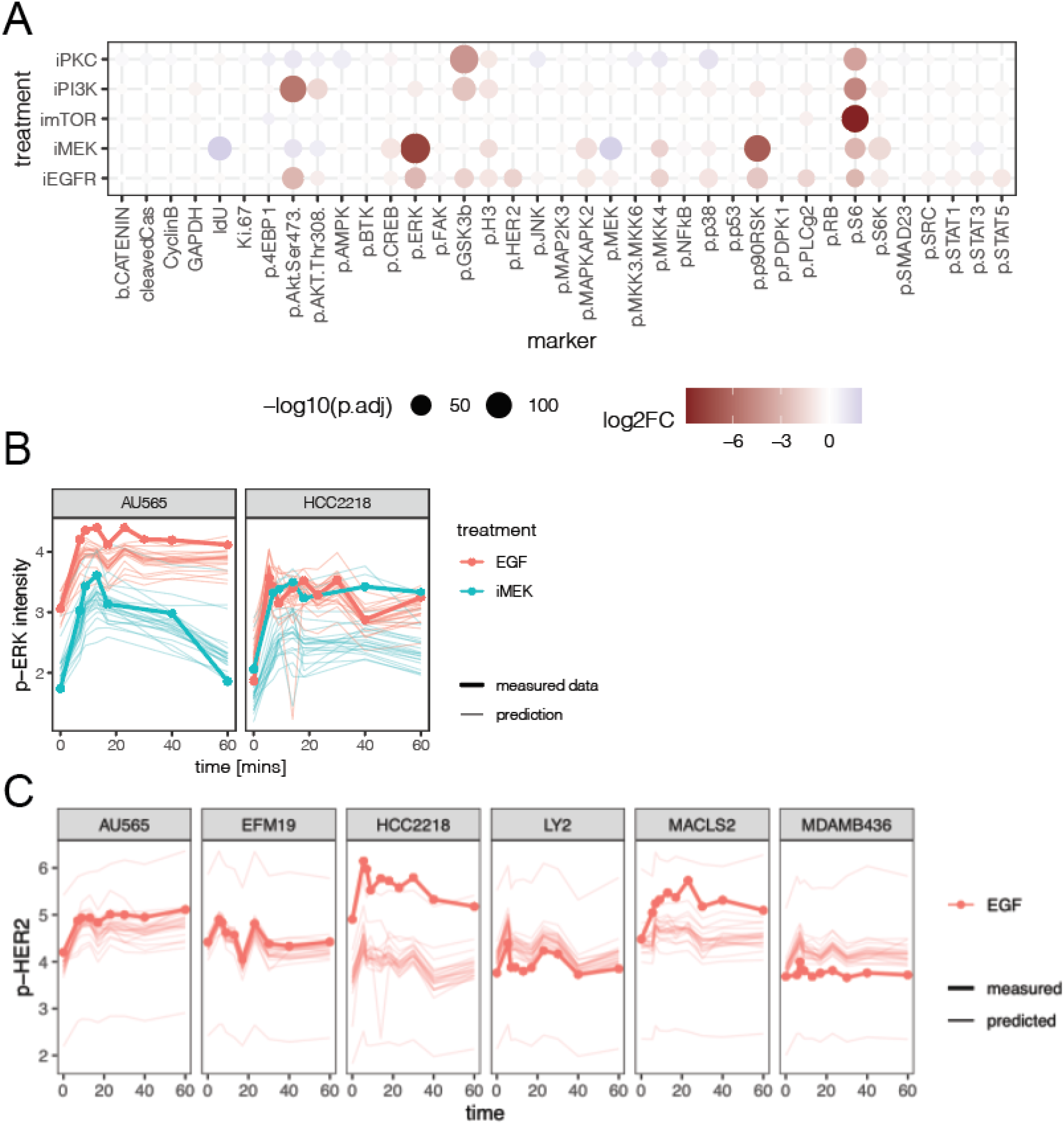
Effect of kinase inhibitors. A) The population level effect of inhibitors on the phosphosites averaged across all 67 cell lines. (log_2_ fold change of signal level between cells treated with EGF and EGF in combination with a kinase inhibitor). B) Measured and predicted population level p-ERK response to EGF and EGF + iMEK inhibitor. C) Measured and predicted population level p-HER2 response to EGF and EGF + iEGFR inhibitor.

The estimates of p-HER2 in HCC2218 cells also showed a large prediction error (Figure 3C). This is mostly due to the large shift in the initial (time 0) signal between the predictions and the data; HCC2218 shows higher activation of p-HER2 than other cell lines in the test set. This also emphasizes the importance to measure and incorporate the basal phosphorylation of predicted markers in all cell lines in the modelling whenever possible.

In summary, this subchallenge shows that models can learn the signaling relationship of the markers on a single cell level and they can predict the response of individual markers. Further, the basal activation of the maker in the new cell line can improve these predictions and particular attention has to be paid on non-canonical signaling effects.

### 2.2 The missing conditions subchallenges

In SC2 and SC3 we challenged participants to predict the effect of known and new inhibitors at the single cell level. We asked them to estimate the entire single cell response of selected cell lines to specific kinase inhibitors, i.e. all markers at all measured time points. This is a fundamentally different task from SC1, because all the nodes are unknown in the predicted condition and therefore the models have to capture the signaling relationships of all the markers to predict between conditions (Figure 1).

The task in SC2 was to predict the response to EGF stimulus in combination with four inhibitors (targeting EGFR, MEK, PI3K and PKC (Supp. Table 3)) in twelve test cell lines (Supp. Figure 2). The training data in SC2 contained the response of the other cell lines to these inhibitors, as well as the basal phosphorylation data for those cell lines. In contrast, in SC3 we asked the participants to predict the effect of an mTOR inhibitor, without giving them any training data for this specific condition. We only provided the name of the inhibitor and the concentration applied in the experiments (Supp. Table 3), and data upon treatment with other inhibitors on those cell lines. Since the effect of this inhibitor could not be observed from training data, participants were encouraged to use prior knowledge for this task.

The predictive models were scored and ranked based on 10,000 single cell predictions submitted for each target cell line and condition by each team. We scored the predictions using population level statistics; the mean value of each marker and the covariance between marker pairs (see Methods) in each condition. This way we capture not only the accuracy of individual markers, but also the changing interplay (covariance) between markers in the score (Figure 4C). A reference model was built using a random forest predictor (see Methods).

**Figure 4:**
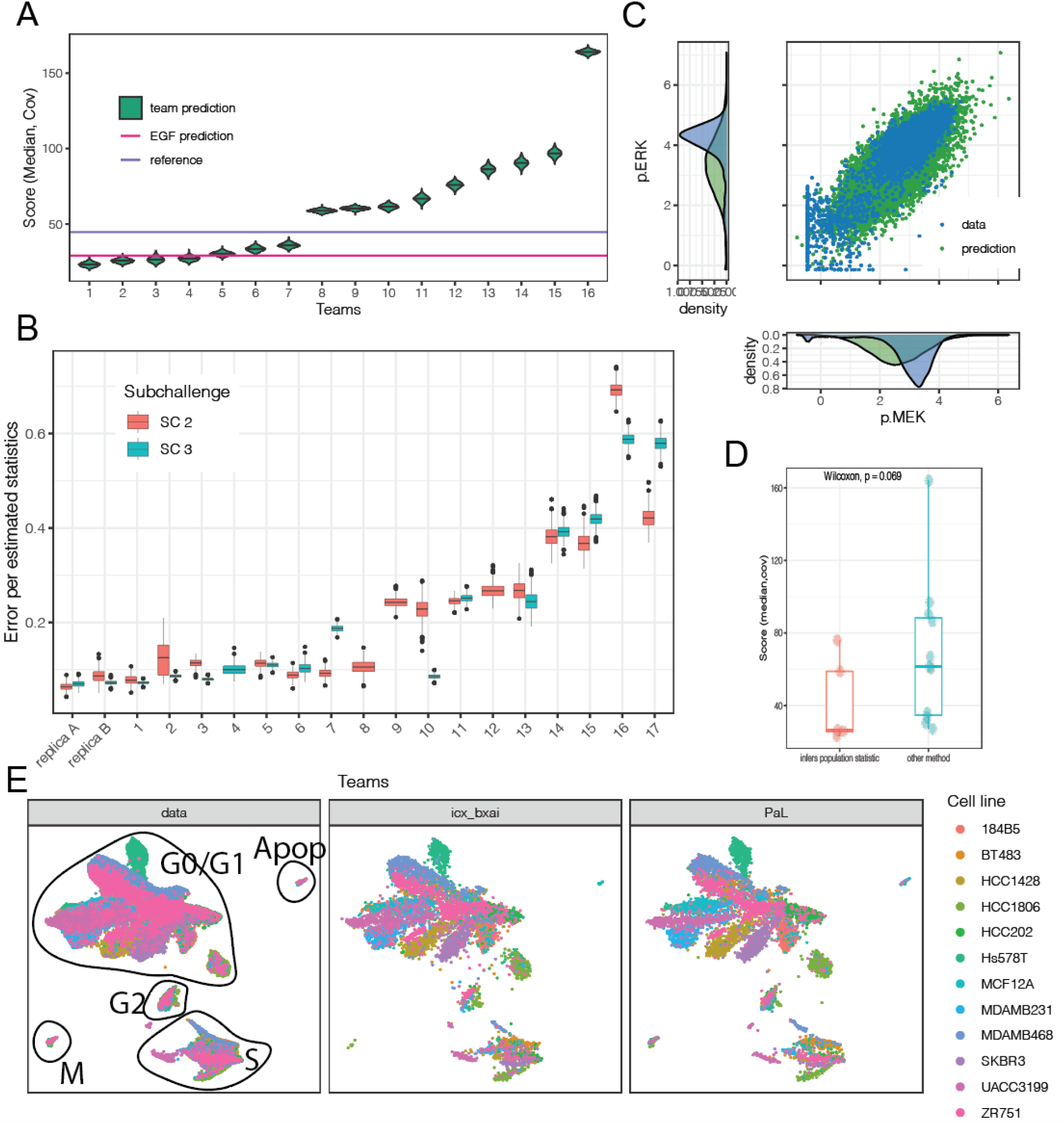
Summary of subchallenge 2 and 3. A) ranking of the teams together with the score of reference predictions in SC2 (see text). B) Comparison of teams’ predictions for SC2 and SC3; the distribution of prediction error is obtained via boostrapping. In addition, the prediction error that one would obtain using either replica A or replica B as predictions is shown on the subset of data that overlaps between both replica. C) Comparison of single cell predictions with the data for the test cell line MDAMB468, iEGFR perturbation, 7 minutes after treatment. D) comparison of performance of teams using statistical inference and then applying Gaussian sampler against teams using other methods. E) Test data shown on two-dimensional Uniform Manifold Approximation and Projection (UMAP) with annotated clusters that belong to different cell-cycle phases (left); projection of the challenge winner’s prediction on the same manifold (middle), projection of the fourth team’s, sampling-based prediction on the same manifold (right)

#### 2.2.1 Quality of predictions

As for SC1, we ranked the participants based on bootstrap samples and compared the ranked predictions in SC2 to a reference model that we built before the challenge (Figure 4A, model described in Methods). Seven out of the 16 teams performed better than our reference model (RMSE_reference_ = 44.67), with up to 48% improvement (median improvement: 39.3%). Of note, a predictor corresponding to the lack of effect of the kinase inhibitor, thus using the measurements to the stimulus alone, leads to a score of RMSE_EGF_ = 29.156. This relatively good score is due to the fact that the kinase inhibitors are very specific and influence only a small number of nodes. Surprisingly, only the top four teams managed to perform better.

A subset of the test data (all cell lines, 15 mins after incubation with inhibitor, without EGF stimulation), contains two biological replicates (replica A and B). Their variation can be considered as an estimate of the upper-bound of performance as it corresponds to the experimental error. Therefore, we also scored the participants on this subset of conditions and compared their scores to the variation of the measured individual biological replicas. Since SC2 and SC3 used different numbers of cell lines (12 and 38, respectively), we computed the prediction error per total number of predicted statistics. This allowed us to summarize the two subchallenges together (Figure 4B), however they remain not directly comparable as the test cell lines and conditions were different. With that in mind, in SC2 the mean score of replica A and B (0.0753) is just slightly better than the top performing team (0.0782), which also holds for SC3, where the average score of the two biological replica is 0.0712 and the best team achieved 0.0725. This shows that the best team could predict single cell response with very good accuracy, close to that of a biological replica.

Similarly to SC1, we found that in both SC2 and SC3 the prediction error fluctuated across cell lines (ANOVA, p-value = 8.8e-4), less so across treatments (p-value = 0.034) and not significantly across time (p-value 0.54), (Supp. Figure 5A). The differences across cell lines were more prominent when only the top four teams, which performed better than the reference model, were considered (Supp. Figure 5B).

### Modelling approaches

Interestingly, five teams, including the top three (*icx_bxai, pqui* and *orangeballs*) followed similar modelling approaches for SC2. These teams inferred the median phosphorylation level of the proteins and their covariance matrices for each sample (cell line, treatment and time point) using linear models (Elastic-net by *icx_bxai* and linear regression by *pqui* and *orangeballs*). Further, all three predicted the cells in the target conditions using the available conditions of the same cell line, rather than from the same conditions of other cell lines. In the next step, they sampled from a multidimensional Gaussian distribution that followed the inferred statistics in each condition which resulted in the single cell predictions. Despite all these similarities, the Z-score transformation, handling the missing time points and using other statistical features of the single cell distributions (means, medians and quantiles) led *icx_bxai* to the first place.

We also evaluated multiple scenarios for combining the predictions for SC2 (see Methods). The combination of top N teams led to 1 - 4.2% improvement over the top ranked team for the range of N = 2-8 teams, the best score was achieved with the top four teams (Supp. Figure 6). A random forest model trained on the predictions errors also performed similarly. The combination of any number of random teams occasionally achieved good scores and had a decreasing trend as the number of teams increased, but on average performed worse than simply assuming no inhibition effect.

#### 2.2.3 Intra-cell line heterogeneity

We then analyzed, for SC2, how the different methods generated heterogeneity and how this matched the heterogeneity found in the data (Figure 4E). We could distinguish two fundamentally different prediction methods. Five teams (including the top three teams) used methods that involve the estimation of population level statistics of the cells (mean and covariance) and then used a Gaussian sampler that generated single cells that followed the estimated statistics. Due to the Gaussian sampler, these approaches inherently assume homogeneity. These types of methods showed good performance: teams using this approach achieved on average 38.7% better score (Wilcoxon test, p-value: 0.069) than teams relying on other methods (Figure 4D). Another popular method (3 teams) was to resample and then optionally scale the available training data (e.g. the 4th best team). This method does not rely on a similar assumption of normality, and thus manages to keep the underlying heterogeneity of the cells.

The measured cells show heterogeneity: the single cell measurements have the tendency of forming multiple clusters that are driven by the cell cycle states (see Methods), and where all the cell lines are represented in each main cluster (Figure 4E). The global layout of the predictions that are either based on Gaussian sampler (Figure 4E middle) or based on resampling and scaling real measurements (Figure 4E right) are in good agreement with the measurements (Figure 4E left). However, when we focused on cells in a single cell line and single treatment conditions (Supp. Figure 7), the heterogeneity due to cell cycle states were only visible in the resampling method because resampling inherits the heterogeneity of the sampled cell lines, while the Gaussian sampler based method predicted homogeneously distributed cells.

In summary, the participants predicted the time response of all markers in response to kinase inhibitors, first (SC2) to inhibitors with known effects and then (SC3) to an inhibitor which was absent from the training data. The results show that predicting distributions of single-cells’ populations after perturbations with kinase inhibitors is almost as good as a biological replica and that, at least for some cases, this is possible even if no data is available upon perturbation with those specific inhibitors. The four best teams were better than the reference model and the top team achieved an accuracy similar to a biological replica. Although overall machine learning-based methods performed better, sampling methods were better able at capturing single cell heterogeneity.

### 2.3 The time-course prediction subchallenge

Perturbation data contains invaluable information about the directionality, magnitude and timescale of the cell response. It also reveals causal interactions of nodes, therefore it is often used for network inference (Zoppoli, Morganella, and Ceccarelli 2010). However, do we need to do perturbation experiments on each and every cell line or is this information already available from the genome, basal protein expression and basal phosphorylation? We formulated subchallenge 4 (SC4) to investigate how well the dynamic response of cell lines can be predicted from their unperturbed, basal state. The participants had to predict the population level response (median of single cells) of five test cell lines to EGF stimulus alone and in combination to four kinase inhibitors across the measured time course (Supp. Figure 2). As in previous subchallenges, they were allowed to build their models based on the training cell lines data (for some of which data under all conditions is available) and use the basal omics (unperturbed, basal proteomics and phopshoproteimics and genomics) data of the test cell lines.

#### 2.3.1 Quality of the predictions compared to an average cell line

We ranked the predictions of the participants based on the RMSE (Figure 5A) and compared them with a reference prediction (model built before the challenge, see details in Methods) and with a baseline model in which we simply averaged all the training cell lines in each condition. Clearly, the baseline model captures the general trends (Figure 5D) and does not capture any cell line specific responses. Based on the scoring metric (RMSE), 11 of the 20 teams predicted better than the baseline model and the top team performed 40% better than the baseline (score of the baseline: 0.428, reference model: 0.3408, top team: 0.260).

**Figure 5:**
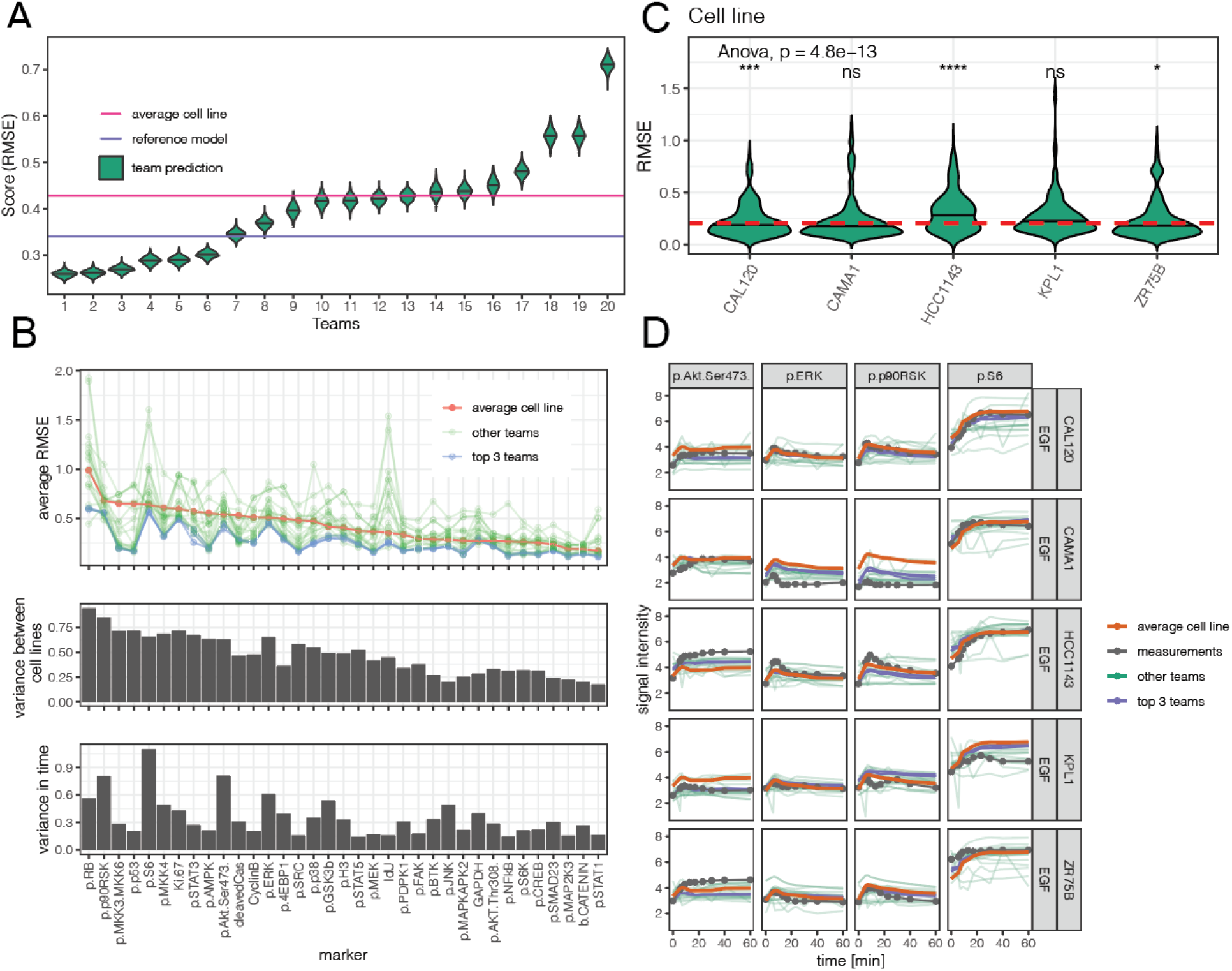
Prediction of dynamic response from basal omics. A) ranking of teams in SC4 based on RMSE with a reference of the averaged training cell lines. B) comparison of the error of predictions and mean of the training cell lines for each marker (top), variance of each marker between cell lines and mean variance of each marker in time. C) Comparison of prediction errors in each cell line. D) time-course comparison of the dynamic response of the test cell lines with the average of the training cell lines and the teams’ predictions (highlighting the top 3 teams).

The top three participants outperformed the average cell line model (baseline model) for all the markers (Figure 5B) except for *GAPHD*, for which their prediction error was similar to the average cell line model. Comparing the predictions with the baseline model, the RMSE was not the same for all markers: certain markers, for example *p*.*p53* and *p*.*MKK3/6*, were predicted much better by the top teams than by the average cell line model (Figure 5B).

To understand which markers can be modelled more accurately, we compared the marker variance across cell lines to the prediction errors (Figure 5B) and we found that the variance correlates strongly with the error in the average cell line (corr. coeff: 0.944), but less to the errors in the participants’ prediction (corr.coeff: 0.73). This means that the participants’ models performed better than the average cell line model, especially for the markers that changed strongly between cell lines.

Further, we also compared the prediction errors with the longitudinal variance of the markers (Figure 5B) that tells us how strongly the markers respond to the perturbation in time. The longitudinal variance correlated more with the participants error (corr. coeff: 0.858) than with the average cell line prediction error (corr. coeff: 0.501). We can also see that the participant’ prediction were much better than the average cell line model when the marker longitudinal variance was lower (Figure 5B), i.e. when the marker changed less in time.

Surprisingly, in contrast to SC1, the prediction errors were similar across cell lines. Although the distribution of the prediction error across the test cell lines showed statistically significant differences (ANOVA test, p-value = 4.8e-13, Fig 5C), the differences are not strong: the largest difference in RMSE (0.0528) was found between cell line HCC1143 and the mean cell line. This shows that modelling approaches generalises well across cell lines.

In summary, in the last subchallenge, the goal was to predict the dynamics of the population response to perturbations from unperturbed basal data. Only 11 teams achieved better performance than random, underscoring the difficulty of the task. Unsurprisingly, predictions were particularly good for markers that did not change across time, but changed across cell lines.

## 3 Summary and Discussion

New technologies allow us to interrogate cell-signaling at the single-cell level. New types of data come with new challenges for data analysis and it is crucial to understand and tackle the limitations of existing computational methods. Towards this goal, we organised the Single Cell Signaling in Breast Cancer DREAM challenge to draw the attention of the computational community to single cell prediction problems, obtain state of the art solutions, and evaluate their performance. Our results provide a baseline for further development and a reference for the community to learn what approaches work best and what the limitations are.

The participants built predictive models from a rich and complex data set, composed of the largest single cell signaling dataset to date and complementary bulk data types. In Subchallenge 1, the majority of the teams predicted missing nodes in the signaling network building models based on all the other measured nodes in each single cell and a summary (e.g. principal components) of the RNAseq and proteomics data. These types of models were trained on the cell lines provided and then used to predict the test cell lines. This implicitly assumed that the relationship among the nodes can be transferred between cell lines. The predictions strongly correlated with the test data, and explained large amounts of variance, suggesting that there are indeed signaling interactions transferable among cell lines.

However, we found two major pitfalls in the predictions. First, the models captured much better the trends than the absolute value of nodes’ activities in cases where the basal marker intensity of the test cell lines were much higher than of the training cell lines. If the absolute values are important, this can be an important limitation, but not for downstream analysis where the data is rescaled (e.g. (Tognetti et al. 2020)). In addition, the functional effect of at least some main pathways is driven by fold-changes, and thus predicting these, even if not being able to do so accurately for the absolute values, is helpful (Adler and Alon 2018). The second pitfall is related to rare signaling cases: in the majority of cell lines MEK phosphorylated ERK and the inhibition of MEK resulted in the lower phosphorylation of ERK. However, in a few cell lines, ERK was phosphorylated independently of MEK. This phenomenon was not captured by the predictions either because the number of such cell lines were low in the training data or because there was no information on this alternative relationship in the data, for example, if this phenomenon depends on the expression of a non-measured node.

The approach to predict the whole signaling response of the cells to kinase inhibitors (Subchallenges 2 and 3) was fundamentally different from Subchallenge 1. The top performing models learnt the signaling relationships among the nodes and how these changed between conditions within each cell line, and transferred this knowledge to the target condition. Further, instead of predicting each individual single cell directly, the top performing models derived statistical descriptors of the markers in each condition, predicted the distribution of cells in the target condition and then sampled that distribution to generate the single cell predictions. We found that the predictions of the top team were almost as accurate as biological replicas. Although models that inferred directly the median and variance of cells performed better than models based on resampling similar cell lines, the single cell analysis showed that the latter approach performed better at capturing the cell lines’ heterogeneity.

In Subchallenge 4 the task was even more challenging: predict the time course response of cell lines for which only basal (before perturbation) data was provided. Many methods performed better than using an average of the provided cell lines as a predictor, showing that participants were able to capture the differences between the cell lines, particularly for phospho-proteins that did not change much across time, but varied between cell lines.

Across all subchallenges, we found that the ensemble of predictions of multiple teams is better than an average team, which means that the risk for a bad prediction can be mitigated by combining multiple approaches. However, the combined prediction is not significantly better than the best team, which is partially due to the fact that teams used relatively similar approaches, and also the best team already used an ensemble of models.

The results of our challenge also provide general guidelines to design experiments aimed to characterize signaling events at the single-cell level. In a setup where each data point has essentially the same cost, it seems that one could reduce the time resolution, as predictions across time points were particularly effective. In addition, given the broadly good results from Subchallenge 1, one could design experiments where only subsets of markers are measured in the different conditions, as long as each phospho-protein is measured in at least one condition for each cell line, even if some of these are bulk measurements.

Further challenges could also address how to predict specifically the basal single-cell signaling from other omics data. If this is achieved, one could predict the single-cell dynamics of signaling networks directly from bulk omics data.

In summary, the results of this DREAM challenge provide a snapshot of our collective ability to predict signal transduction at the single-cell level. The best performing methods can be applied to other datasets, and used as a baseline for those interested in developing methods for these aims. To support those further developments, all data and descriptions of methods are made freely available for the community to use.

## 4 Methods

### 4.1 Single Cell Signaling in Breast Cancer DREAM challenge

The challenge was divided into three rounds which lasted between two to four weeks each. In each round participants were limited to maximum three submissions, to avoid learning on the test dataset. After each round, the current scores were publicly presented on a scoring board. In total 127, 77, 61 and 83 valid predictions were submitted by 22, 16, 14 and 25 teams for each subchallenge, respectively.

A requirement for the challenge participation is a summary of method, which is publicly available (https://www.synapse.org/#!Synapse:syn20366914/wiki/593925) and open source code that reproduce the results. Further, we collected information on the methods via a questionnaire (Supp. File 1).

### 4.2 Scores and robust ranking by resampling

In subchallenge 1, we computed the root mean square error between the measured data *y* and prediction *ŷ* for each marker *m*, separately in each condition, on single cell level. In a particular condition, there are *N*_*cl,tr,t*_ single cells that are measured and predicted in each test cell line *cl*, treatment *tr*and time point *t* after the perturbation. The RMSE is computed as

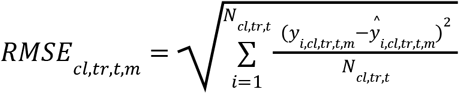

For the final score of a team we computed the average RMSE error across the five predicted markers (p-ERK, p-PLCg2, p-HER2, p-S6, p-AKT_S473), six cell lines (AU565, EFM19, HCC2218, LY2, MACLS2, MDAMB436), and all measured timepoints (11 in stimulated experiments, 7 in case of combined stimulus and inhibition):

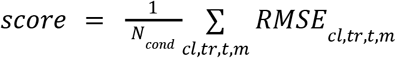

We found that the RMSE score changes in each condition, most significantly based on the cell line that was predicted. Therefore, for a robust ranking of participants, we resampled 1000 times the conditions used to compute the final score with replacement and computed the bootstrap score on each sample. Finally, the team are compared by Bayes factors based on the bootstrap scores:

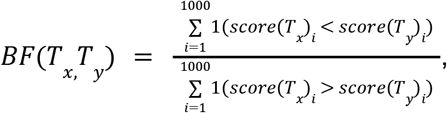

where *score*(*T*_*x*_)is the score of team *T*_*x*_, 1() is the indicator function.

In subchallenges 2 and 3 the participants submitted 10’000 representative single cell predictions (predicting *N*_*marker*_ = 35 markers) for each validation cell line and condition (treatment and time). For each condition *i*, we compared the mean marker value from the gold standard data (µ)with the mean of the predicted sample 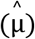

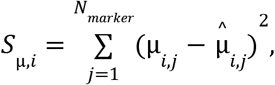

and also for the covariance of marker pairs:

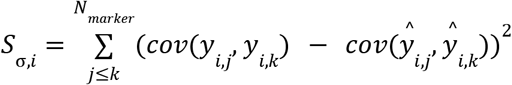

and combined the two parts equally when computed the average across all cell lines and conditions (*N*):

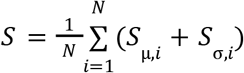

For a robust ranking, we created bootstrap samples of scores by resampling the conditions and cell lines with replacement. Then we computed the Bayes factors between teams, based on the bootstrap scores.

In subchallenge 4 the participants predict the population mean response to perturbation over time for each maker. Therefore, for each cell lines and treatment we computed the RMSE across time:

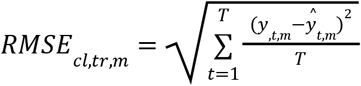

The final score is computed by averaging the error for all conditions.

For a robust ranking, we created bootstrap samples of scores by resampling the treatments and cell lines with replacement, as before. Then we computed the Bayes factors between teams.

### 4.3 Reference models

In SC1, our model predicted the missing marker based on the measured markers in each individual cell. We built a random forest model for each marker independently, across all conditions using the *ranger* R package(Wright and Ziegler 2017). The model features included all the measured markers and time as continuous variables, and the treatments, starvation status (cells in treatment “full” were not starved) and stimulated status (cells at time 0 and in “full” treatment are not stimulated with EGF) as one-hot-encoded variables. The model was trained on a subset of the training cell lines: we selected 7 cell lines randomly (*BT474, CAL148, HBL100, MCF7, MDAMB157, TD47D* and *ZR7530*) and 500 random cells from each condition.

The idea for the reference model in SC2 was to predict the means and covariance matrix of the markers in the missing conditions based on the available conditions, and then use a simulator that generates samples from a multivariate normal distribution following the estimated mean and covariance values. Thus, first we computed the mean and covariance matrix of the markers in each cell line, treatment and time point. Then we built an independent random forest model for each statistical variable *y* (mean of a marker or entry in the covariance matrix), for each cell line and time point:

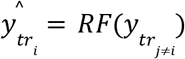

For example, the mean value of marker *Ki-67*, in cell line *BT474*, time 0, treatment iMEK was predicted based on the available values of *Ki-67* levels in the same cell line, in time 0, in {EGF, iEGFR, iPI3K, iPKC} conditions. For the training of the models we selected the same 7 training cell lines as in SC1.

We haven’t built a reference model for subchallenge 3.

In subchallenge 4, we hypothesized that each marker response (*y*_*cl, tr, time*_) can be described as the superposition of the average cell line response 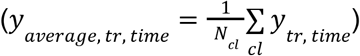 and cell line specific response difference (*d*_*cl,tr,time*_), i.e.

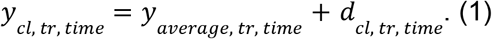

The average cell line response (*y*_*average, tr, time*_) was calculated by taking the mean of all the training cell lines for each marker, in each condition (treatment and time point). Note that all the three terms in Eq. (1) can be easily calculated for the training data. Then we built a random forest model that predicts the cell line specific response (*d*_*cl, tr, time*_) based on the data that is available for the test conditions, i.e. based on the proteomics data and based on the basal activation of markers (“full” condition). We also used the treatment, time and the stimulation status (i.e. if time>0) as model features. We trained a model for each reporter independently based on all the training cell lines.

### 4.4 Random and systematic combinations of predictions

Predictions of randomly selected teams and the top performing teams were aggregated. Different sizes of ensembles (n) were scored, from the minimum of one team up to the maximum of all teams. In subchallenge 1 and 4 the predicted values of different teams were combined by taking the median of the predictions. In subchallenge 2 and 3 representative cells were predicted and scored based on the statistics. Here we used two approaches: (1) predictions were combined by sampling an equal number of predicted values per condition from all selected submissions and (2) computing the statistics for each submission and then averaging across the selected teams. In all subchallenges the resulting ensembles were scored the same way as individual submissions. For the random combination of teams, we repeated the procedure 100 times to see the distribution of score.

### 4.5 Machine learning based combination of predictions

We used Random Forest (RF) models in SC1 and SC2 to combine the predictions for the test cell lines. We used a cross-validation scheme for the model training and for the predictions to avoid overfitting these RF models and to generate predictions for all the test cell lines. We divided the data into six folds: in SC1 each fold of data contained a single test cell line and in SC2 each fold contained two cell lines. This way the training and predictions of the RF models also mimic the setup of the subchallenges, i.e. they are trained on part of the cell lines and predict other cell lines. Then in six iterations, we trained our RF models on five folds and predicted the sixth fold until we had predictions for all the test data and could compare the performance of RF predictions with the participants’ predictions.

In SC1 we trained a random forest model per marker and used the single cell predictions from the teams and the time point as continuous variables and the treatment and predicted marker as one-hot encoded features. Due to the chosen cross validation scheme we could not use “cell line” as a feature. Therefore, we also included the 32 non-predicted marker values as model features.

In SC2, our RF predicted the mean and covariances per condition using the mean and covariance matrix from the participants predictions. The model also included as features the median expression per marker at treatment EGF. The predicted statistics were used to create a multivariate normal distribution, from which 10’000 cells per condition were sampled and subsequently scored.

### 4.6 Cell cycle phases

Cell cycle phases were identified from the single cell data based on the markers *Histone H3* (p.H3), *iododeoxyuridine* (IdU), *retinoblastoma protein* (p.RB) and *cyclinB* (Supp. Figure 8). Following *(Behbehani et al. 2012)*, G0- and G1-phase are both characterised by low expression of IdU, p.H3 and cyclinB, but we could not differentiate between G0 and G1 based on the expression of *retinoblastoma protein* (p.RB). High expression of IdU and low expression of p.H3 characterizes S-phase; high expression of p.H3 with low IdU identifies M-phase; finally, low expression of IdU and p.H3 with high expression of cyclinB identifies G2 phase. Apoptotic cells were detected by high cleaved *Caspase 3* (cleavedCas) levels.

### 4.7 Code availability

All the challenge data, predictions, code and write-ups for the predictions are available at https://www.synapse.org/singlecellproteomics. The code and data to reproduce the results presented in this manuscript is available as a CodeOcean capsule, Single Cell Signaling in Breast Cancer DREAM challenge analysis (https://codeocean.com/capsule/7326564/tree), DOI: 10.24433/CO.6078101.v1.

## Acknowledgements

B.B. was supported by a SNSF R’Equip grant, a SNSF Assistant Professorship grant, the SystemsX Transfer Project “Friends and Foes”, the SystemX grants Metastasix and PhosphoNEtX, a NIH grant (UC4 DK108132), the CRUK IMAXT Grand Challenge, and by the European Research Council (ERC) under the European Union’s Seventh Framework Program (FP/2007-2013)/ERC Grant Agreement n. 336921

Thanks to Natalie de Souza and Olga Ivanova for feedback on the manuscript. Further thanks for Ricardo O. Ramirez-Flores for his suggestions and discussions during the preparation and evaluation of the challenge and for Pablo Meyer for his feedback on scoring and combined predictions.

## Consortia

Single Cell Signaling in Breast Cancer DREAM Consortium members: Augustinas Prusokas, Alidivinas Prusokas, Renata Retkute, Anand Rajasekar, Karthik Raman, Malvika Sudhakar, Raghunathan Rengaswamy, Edward S.C. Shih, Min-jeong Kim, Changje Cho, Dohyang Kim, Hyeju Oh, Jinseub Hwang, Kim Jongtae, Yeongeun Nam, Sanghoo Yoon, Taeyong Kwon, Kyeongjun Lee, Sarika Chaudhary, Nehal Sharma, Shreya Bande, Gao Gao fan zhu Cankut Cubuk, Pelin Gundogdu, Joaquin Dopazo, Kinza Rian, Carlos Loucera, Matias M. Falco, Martin Garrido-Rodriguez, Maria Peña-Chilet, Huiyuan Chen, Gabor Turu, Laszlo Hunyadi, Adam Misak, Baosen Guo, Wencai Cao, He Shen, Lisheng Zhou, Xiaoqing Jiang, Pieta Zhang, Aakansha Rai, Rintu Kutum, Sadhna Rana, Rajgopal Srinivasan, Swatantra Pradhan, James Li, Vladimir Bajic, Christophe Van Neste, Didier Barradas-bautista, Somayah Abdullah Albarade, Igor Nikolskiy, Musalula Sinkala, Duc Tran, Hung Nguyen, Tin Nguyen, Alexander Wu, Benjamin DeMeo, Brian Hie, Rohit Singh, Jiwei Liu, Xueer Chen, Leonor Saiz, Jose M. G. Vilar, Peng Qiu, Akash Gosain, Anjali Dhall, Dinesh Bajaj, Harpreet Kaur, Krishna Bagaria, Mayank Chauhan, Neelam Sharma, Gajendra Raghava, Sumeet Patiyal, Jianye Hao, Jiajie Peng, Shangyi Ning, Yi Ma, Zhongyu Wei, Atte Aalto, Jorge Goncalves, Laurent Mombaerts, Xinnan Dai, Jie Zheng, Piyushkumar Mundra, Fan Xu, Jie Wang, Krishna Kant Singh, Mingyu Lee.

For affiliations, team names and contacts, consult supplementary File 2.

## Authors contributions

AG, MT, JSR and BB conceived and designed the Single Cell Signaling in Breast Cancer DREAM challenge with the help of TY and VC. AG and AD analysed the challenge results with the help of MT and JT, under the supervision of JSR. The top-performing approaches were designed by BG, WC and HS. The DREAM Consortium provided predictions, method implementations and descriptions. AG and JSR drafted the manuscript with inputs from MT, AD and JT. All authors read, commented, and approved the final manuscript.

