## Supplementary material for "Cell-to-cell and type-to-type heterogeneity of signaling networks: Insights from the crowd": Supplemetary material

### 5 Supplementary Materials

Supplementary File 1: questionnaire about the methods in excel.

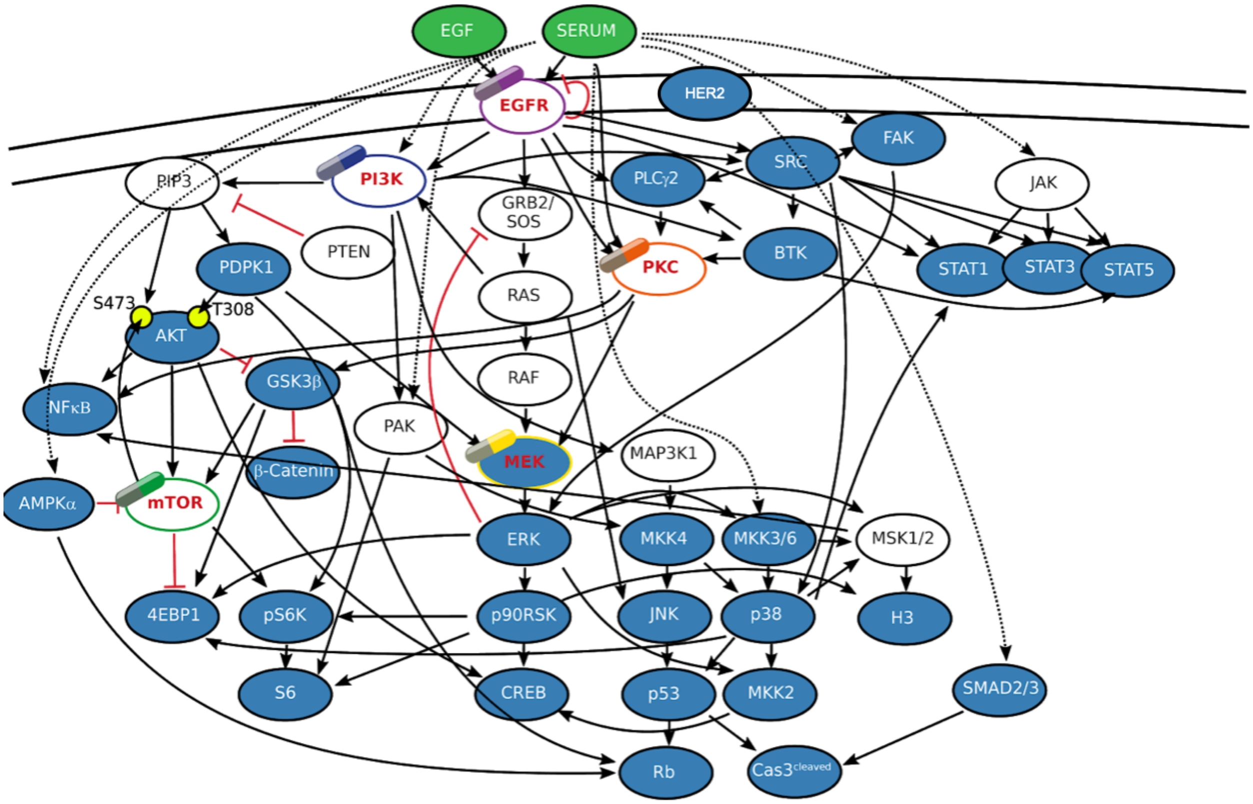

**Supplementary Figure 1: Measured phospho-proteins (in blue) and perturbation targets (indicated by drug symbols) in the single cell experiments.**

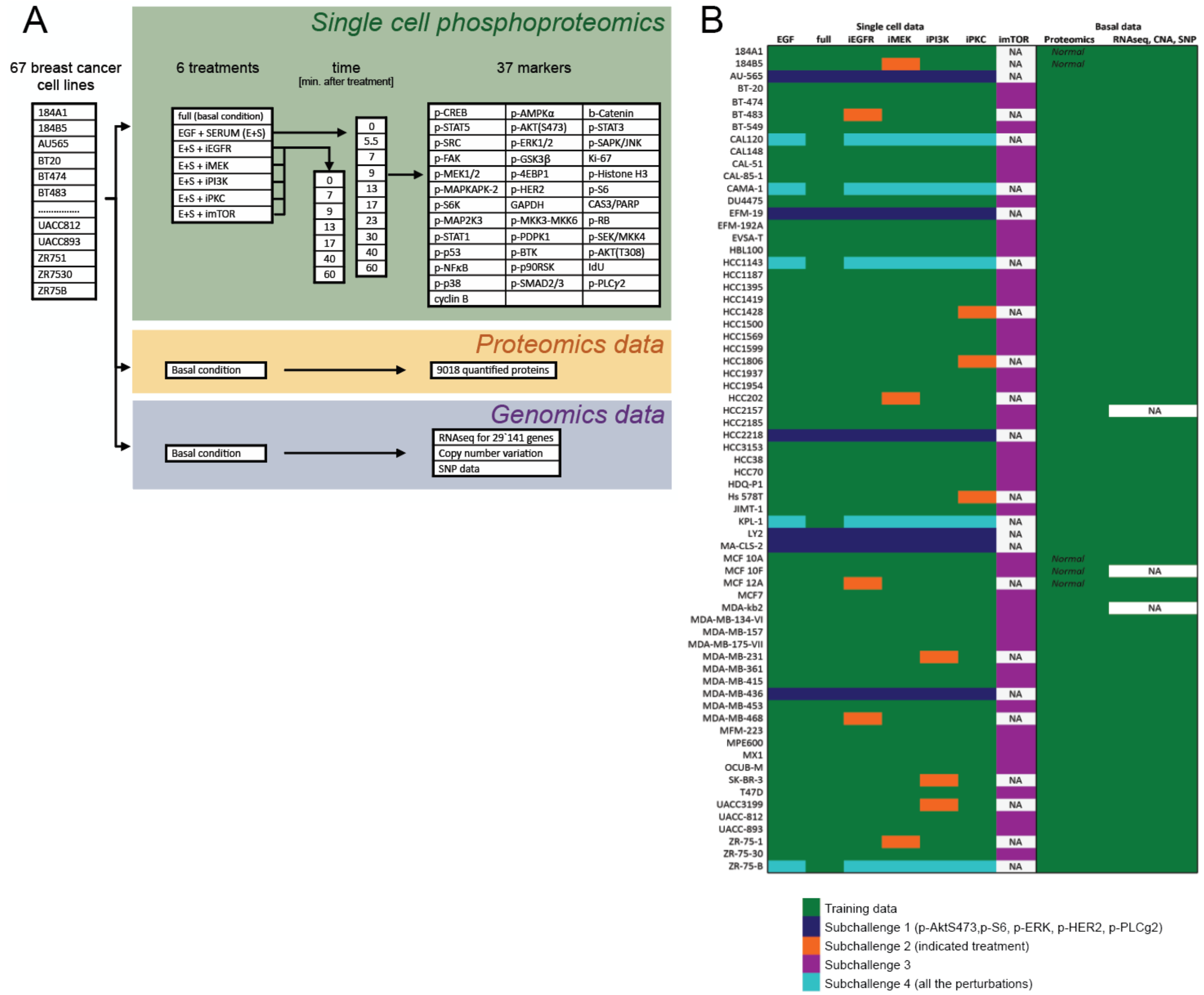

**Supplementary Figure 2. Overview of data availability in the challenge.**
Panel A shows the three different types of data provided for the participants and the experimental conditions available for the single cell phosphorylation data. Panel B shows the distribution of the cell line data between training data (available for participants) and test data (used for scoring). The cells were starved before stimulation with EGF and incubated with the inhibitors for 15 minutes. The initial activation refers to the phosphorylation level at time 0, right before adding the EGF stimulus.This is different from basal activation, which represents the activation of cells growing in normal condition.

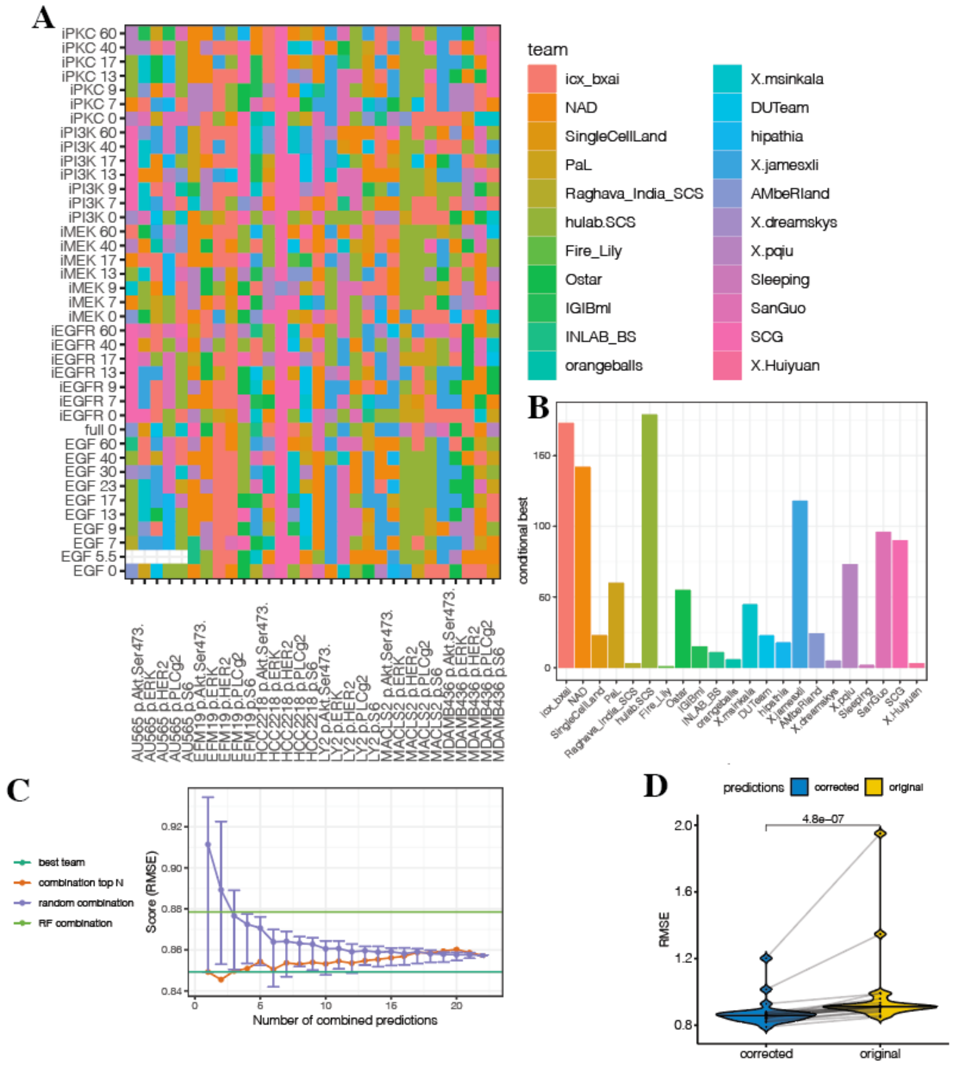

**Supplementary Figure 3. Performance of teams, combined predictions and improved prediction in subchallenge 1.**

*A) the tile plot shows the best performing team for each cell line, condition, treatment and time. B) number of conditions where a team performed the best. C) The performance of combined predictions depending on the number of considered teams (see Methods for details) D) improvement of teams' score after correcting their predictions by the difference between measured and predicted basal phosphorylation in each cell line.*

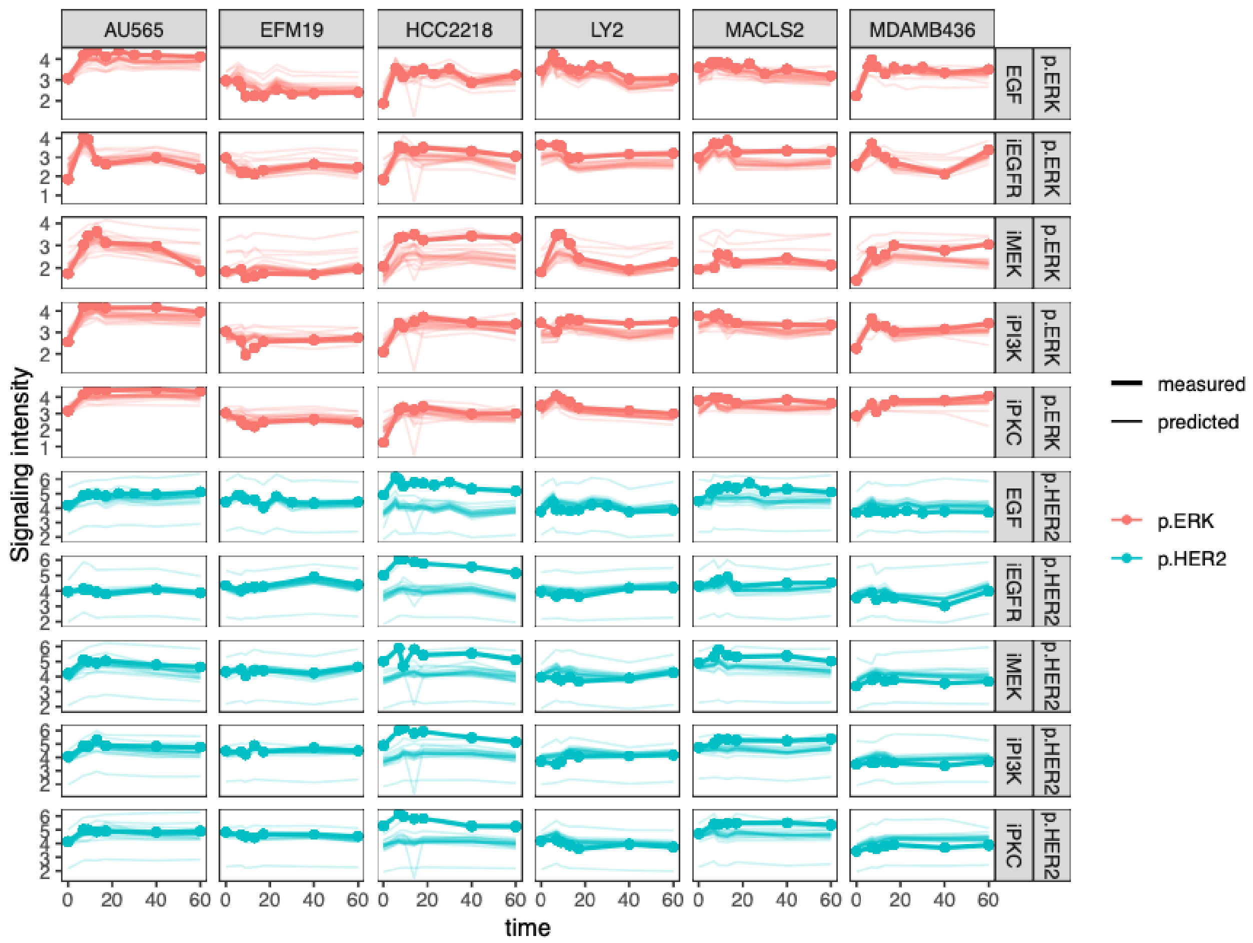

**Supplementary Figure 4 Mean measured data (thick line with dots) and predictions (thin, semi-transparent lines) in SC1.**

*Cell lines respond differently to the same perturbation (in rows). The effect of EGF stimuli in combination with the kinase inhibitors result in different responses. The median predictions of the teams are mostly well overlapping with the measurements. In case of larger differences (e.g. HCC2218, p.HER2, all treatments) the difference between prediction and measurements are almost constant in time.*

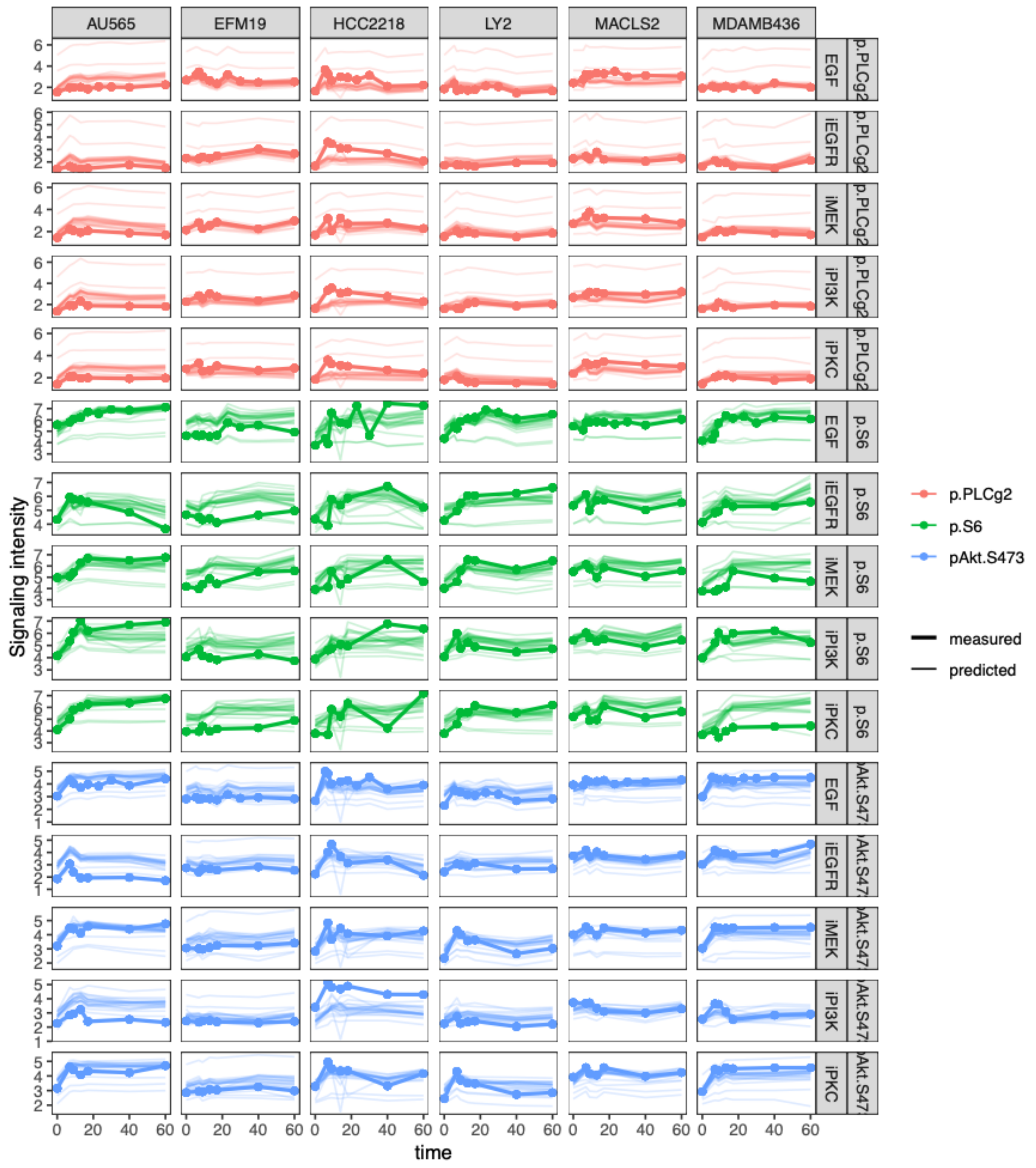

**Supplementary Figure 4(cont.)**

*Mean measured data (thick line with dots) and predictions (thin, semi-transparent lines) in SC1 for the remaining markers.*

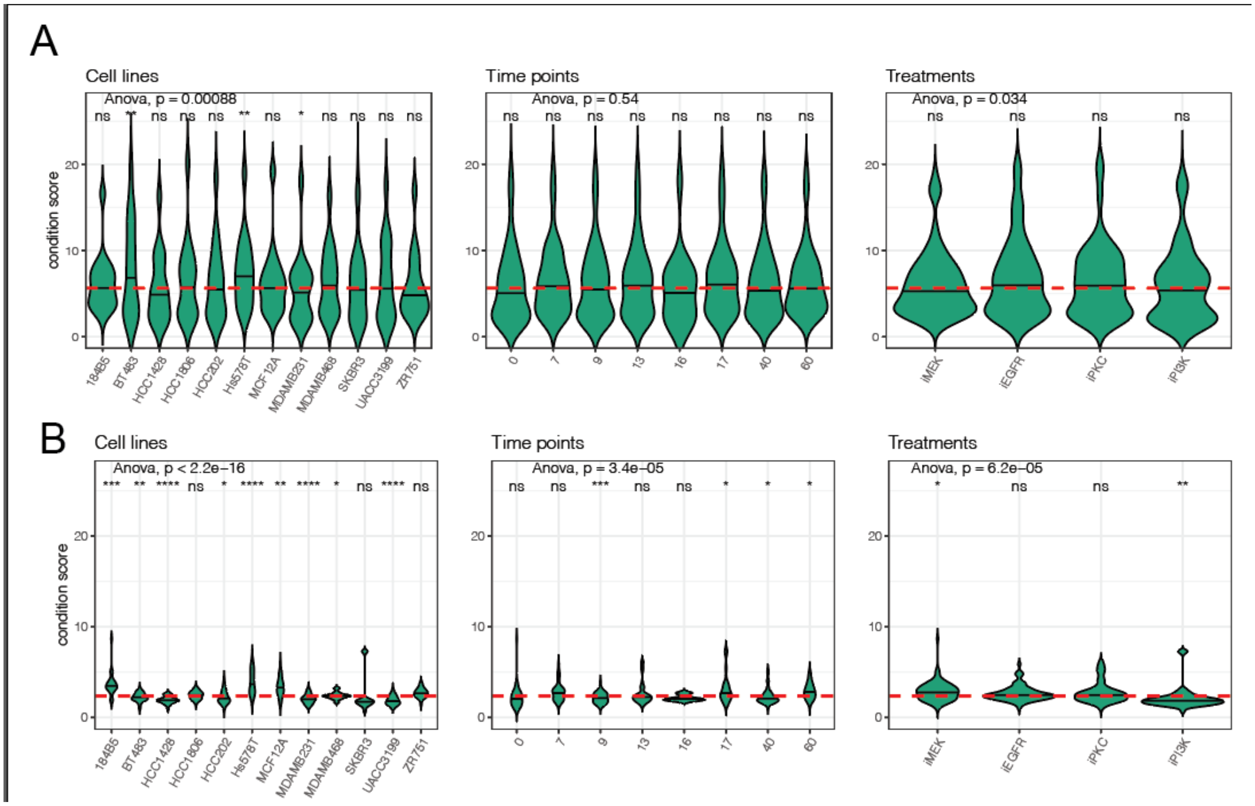

**Supplementary Figure 5**

*distribution of the prediction error score (squared difference of predicted and real median and covariances) across cell lines, treatments and time points in SC2. Each violin contains the predictions of A) all the teams, B) top 4 teams in each condition. The significance of a t-test between the individual groups and the global RMSE is shown above each violin plot ( *: p<0.05, **: p<0.01, ***: p<0.001, ****: p<0.0001, ns: not significant)*

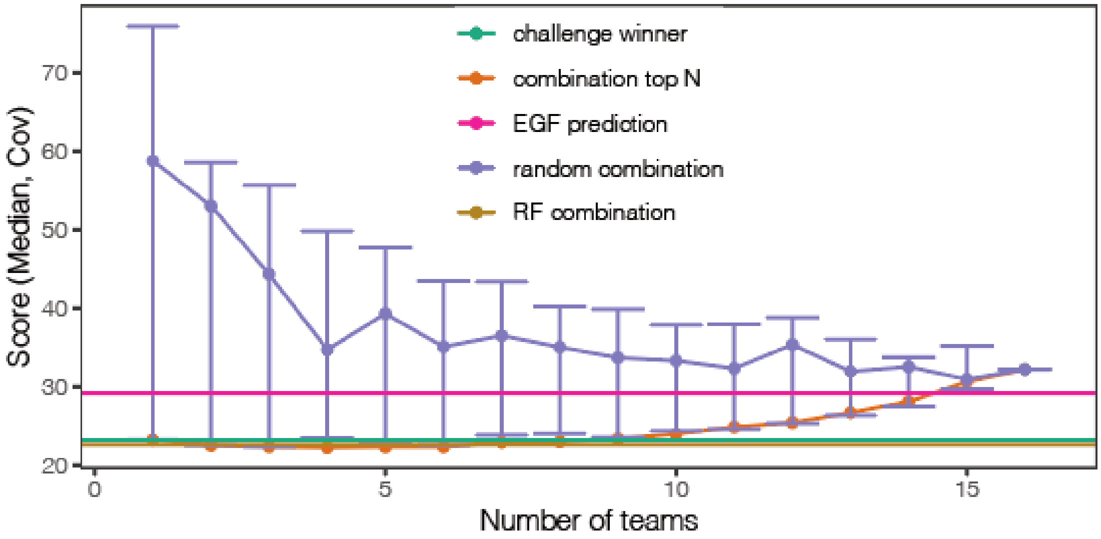

**Supplementary Figure 6. Combination of predictions for SC2.**
*Score of combined predictions based on the number of teams considered.*

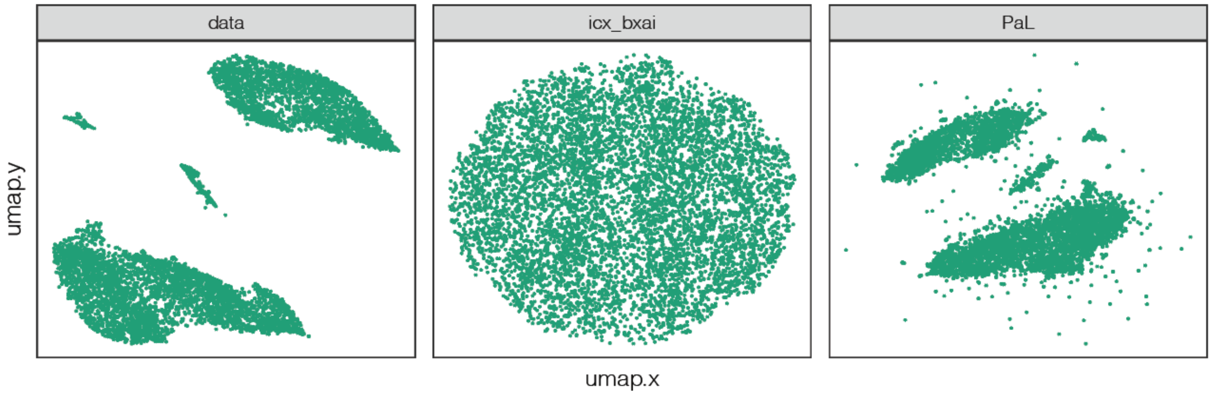

**Supplementary Figure 7.** ***UMAP projection of the data and predictions for the cell line BT483, iEGFR condition in subchallenge 2.*** *The projection of data and predictions independently to a 2D manifold reveals clusters for the data and for the predictions from PaL (used a resampling method), but shows a uniform distribution for the challenge winner's prediction (icx_bxai, used a Gaussian sampler)*

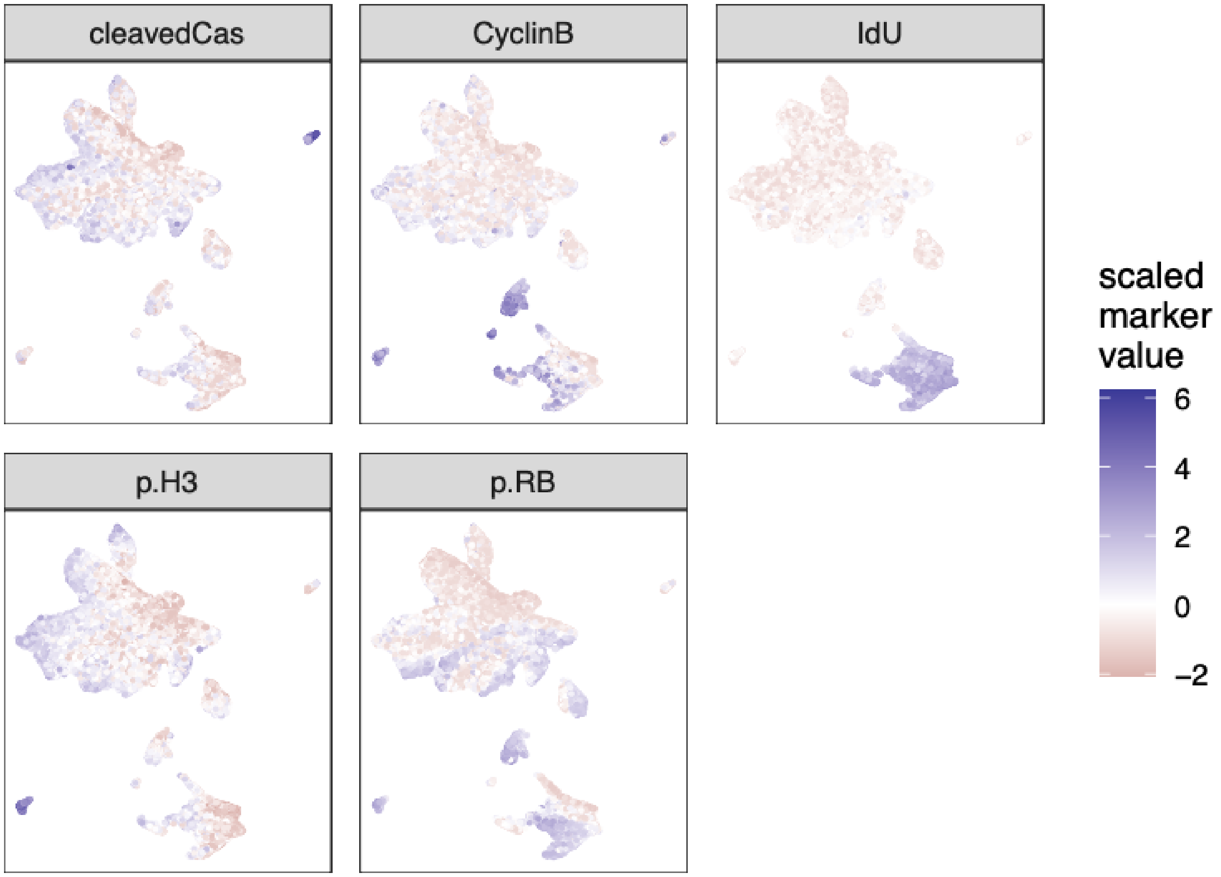

**Supplementary Figure 8**
SC2 test data shown on two-dimensional Uniform Manifold Approximation and Projection (UMAP) colored by markers used to identify cell-cycle phases (see Methods)

### 5.2 Error distributions

We tested in each subchallenges, how the RMSE distributes across cell lines, treatment, marker and treatments. We evaluated this via an ANOVA test, where the null hypothesis is that the RMSE has the same mean value across the variables (Supp. Table 1). Then we compared the RMSE distribution within each group to the global distribution of RMSE (Supp. Table 2).

#### Supplementary Table 1: ANOVA of RMSE in subchallenge 1.

| | Degree of freedom | Sum of Squares | Mean Squares | F-value | Significance | |
|---|---|---|---|---|---|---|
| cell_line | 5 | 14.767 | 2.953 | 179.58 | <2e-16 | *** |
| treatment | 5 | 0.191 | 0.038 | 2.321 | 0.0413 | * |
| time | 9 | 0.269 | 0.03 | 1.817 | 0.0611 | . |
| marker | 4 | 31.335 | 7.834 | 476.342 | <2e-16 | *** |
| Residuals | 1141 | 18.765 | 0.016 | | | |

#### Supplementary Table 2: comparison of RMSE within group versus global distribution via t-test. subchallenge 1

| | | | | | | | | | |
|---|---|---|---|---|---|---|---|---|---|
| SC1: marker vs global | | | | | | | | | |
| | marker | deltaRMSE | groupMeanRMSE | globalMeanRMSE | statistic | p.value | parameter | conf.low | conf.high |
| 1 | p.Akt.Ser473. | -0.0241 | 0.689 | 0.713 | -2.01 | 4.52E-02 | 498 | -0.0477 | -0.000516 |
| 2 | p.ERK | -0.144 | 0.569 | 0.713 | -15.5 | 3.67E-48 | 903 | -0.162 | -0.126 |
| 3 | p.HER2 | -0.142 | 0.571 | 0.713 | -10.6 | 2.48E-23 | 420 | -0.169 | -0.116 |
| 4 | p.PLCg2 | 0.00543 | 0.718 | 0.713 | 0.517 | 6.05E-01 | 654 | -0.0152 | 0.026 |
| 5 | p.S6 | 0.305 | 1.02 | 0.713 | 16.3 | 9.17E-44 | 311 | 0.268 | 0.342 |
| SC1: cell line vs global | | | | | | | | | |
| | cell_line | deltaRMSE | groupMeanRMSE | globalMeanRMSE | statistic | p.value | parameter | conf.low | conf.high |
| 1 | AU565 | -0.0593 | 0.654 | 0.713 | -3.7 | 2.62E-04 | 284 | -0.0909 | -0.0277 |
| 2 | EFM19 | -0.0551 | 0.658 | 0.713 | -3.08 | 2.26E-03 | 268 | -0.0902 | -0.0199 |
| 3 | HCC2218 | 0.246 | 0.959 | 0.713 | 10.8 | 2.82E-22 | 235 | 0.201 | 0.291 |
| 4 | LY2 | -0.0776 | 0.636 | 0.713 | -5.46 | 9.28E-08 | 330 | -0.105 | -0.0496 |
| 5 | MACLS2 | -0.0531 | 0.66 | 0.713 | -4.1 | 5.01E-05 | 372 | -0.0785 | -0.0277 |
| 6 | MDAMB436 | -0.00231 | 0.711 | 0.713 | -0.177 | 8.60E-01 | 367 | -0.028 | 0.0234 |
| SC1: time vs global | | | | | | | | | |
| | time | deltaRMSE | groupMeanRMSE | globalMeanRMSE | statistic | p.value | parameter | conf.low | conf.high |
| 1 | 0 | 0.0369 | 0.75 | 0.713 | 1.87 | 0.0622 | 233 | -0.00189 | 0.0756 |
| 2 | 5.5 | -0.016 | 0.697 | 0.713 | -0.279 | 0.783 | 24.7 | -0.134 | 0.102 |

| | | | | | | | | | |
|---|---|---|---|---|---|---|---|---|---|
| 3 | 7 | -0.000796 | 0.712 | 0.713 | -0.0362 | 0.971 | 183 | -0.0442 | 0.0426 |
| 4 | 9 | -0.00881 | 0.704 | 0.713 | -0.438 | 0.662 | 191 | -0.0485 | 0.0309 |
| 5 | 13 | -0.00173 | 0.711 | 0.713 | -0.0866 | 0.931 | 192 | -0.0412 | 0.0378 |
| 6 | 17 | -0.006 | 0.707 | 0.713 | -0.305 | 0.76 | 194 | -0.0447 | 0.0327 |
| 7 | 23 | -0.037 | 0.676 | 0.713 | -0.884 | 0.384 | 30.7 | -0.122 | 0.0484 |
| 8 | 30 | -0.0244 | 0.689 | 0.713 | -0.537 | 0.595 | 30.4 | -0.117 | 0.0682 |
| 9 | 40 | 0.00479 | 0.718 | 0.713 | 0.238 | 0.812 | 192 | -0.0349 | 0.0445 |
| | 60 | -0.0168 | 0.696 | 0.713 | -0.843 | 0.4 | 193 | -0.056 | 0.0224 |
| SC1: treatment vs global | | | | | | | | | |
| | treatment | deltaRMSE | groupMeanRMSE | globalMeanRMSE | statistic | p.value | parameter | conf.low | conf.high |
| 1 | EGF | -0.0158 | 0.697 | 0.713 | -1.01 | 0.314 | 449 | -0.0465 | 0.015 |
| 2 | full | 0.0463 | 0.759 | 0.713 | 0.971 | 0.339 | 30.3 | -0.051 | 0.144 |
| 3 | iEGFR | 0.00393 | 0.717 | 0.713 | 0.233 | 0.816 | 301 | -0.0293 | 0.0371 |
| 4 | iMEK | 0.0127 | 0.726 | 0.713 | 0.737 | 0.462 | 295 | -0.0212 | 0.0467 |
| 5 | iPI3K | 0.0033 | 0.716 | 0.713 | 0.185 | 0.854 | 288 | -0.0319 | 0.0385 |
| 6 | iPKC | -0.00438 | 0.709 | 0.713 | -0.237 | 0.813 | 282 | -0.0408 | 0.032 |

### 5.3 Experimental perturbations

Supp. Table 3 contains the details of the applied kinase inhibitors and their concentrations.

**Supplementary Table 3: kinase inhibitors applied to single cell perturbation experiments.**

| **Target Molecule** | **Inhibitor Name** | **Alternative Name** | **Supplier** | **Catalog Number** | **Concentration Used** [μM] |
|---|---|---|---|---|---|
| PI3K | GDC-0941 | Pictilisib | LC Laboratories | G-9252 | 0.5 |
| mTOR | Rapamycin | Sirolimus | LC Laboratories | R-5000 | 0.01 |

| | | | | | |
|---|---|---|---|---|---|
| EGFR | Lapatinib | | LC Laboratories | L-4899 | 1.08 |
| PKC | Enzastaurin | LY317615 | LC Laboratories | E-4506 | 3.9 |
| MEK | CI-1040 | PD184352 | Selleckchem | S1020 | 1.7 |
